# Multiparasitism among Schoolchildren of Akonolinga, Nyong et Mfoumou Division, Centre Region of Cameroon

**DOI:** 10.1101/584318

**Authors:** Martin G. Oyono, Leopold G. Lehman, Samuel Fosso, C. F. Bilong Bilong

**Affiliations:** Laboratory of Parasitology and Ecology, Faculty of Sciences, University of Yaoundé I; Parasitology and Entomology Unit, Laboratory of Animal Biology and Physiology, Faculty of Sciences, University of Douala; Laboratory of Parasitology, Mycology and Parasitic Immunology, Yaoundé University Teaching Hospital

**Keywords:** Multiparasitism, Frequency, Determinants, Parasitic association, Schoolchildren

## Abstract

In Sub-Saharan Africa, school-age children are the most vulnerable to parasitic infections and are particularly exposed to multi-parasitism and its potential consequences. This study aimed at determining the intensity of multi-parasitism in Nyong et Mfoumou Division and identifying its determinants. A cross-sectional study took place from September 2017 to July 2018 among pupils of five (05) government schools from the Nyong et Mfoumou Division. Stool samples were collected from each child and examined for protozoan cysts, helminth eggs and larva while blood samples were collected for detection of *Plasmodium* spp and filarial blood stages. In addition, socio-demographic and school environment related information were documented. In total, 416 schoolchildren were recruited; out of which 309 (74.28%) were infected by at least one parasite species. 13 parasite species were found: 03 hemoparasites and 10 intestinal parasites. *Plasmodium falciparum* was the main hemoparasite (37.26%). Amongst intestinal parasites, *Entamoeba coli* were the most common among protozoa (29.33%) and *Ascaris lumbricoides* among helminths (21.39%). The frequency of multi-parasitism was 44.47% and the average species reach was 1.43 ± 0.01 per individual. Four types of multi-parasitism were found (bi-parasitism, tri-parasitism, quadri-parasitism and penta-parasitism); the bi-parasitism (26.68%) was the most common. Significantly statistic associations were found between parasite species such as: *Entamoeba coli*, *Entamoeba histolytica/dispar*, *Ascaris lumbricoides*, *Trichirus trichiura* and *Mansonella perstans* and can generally be explained by the same means of transmission. We conclude that the intensity of multiparasitism among schoolchildren in Nyong et Mfoumou Division is high with predominance in rural areas.

**Author Summary:** Everywhere in Sub-Saharan Africa, school-age children are most vulnerable to parasitic infections, and in higher risk of multiparasitism and its potential consequences. Here, we report results obtained from pupils aged from 4 to 15 years from the Nyong et Mfoumou Division in the Centre Region of Cameroon. Amongst these pupils, 74.28% were infected with one parasite specie at least and 13 parasite species were found in the study area. The highest frequency, 37.26%, was found for *Plasmodium falciparum*. *Entamoeba coli* (29.33%) and *Ascaris lumbricoides* (21.39%) were the most common intestinal protozoa and helminth respectively. The frequency of multiparasitism was 44.47%; 26.68% participants harbored two parasites species concurrently and the maximum number of parasites harbored by one host individual was 5. The risk of multiparasitism was significantly higher for females, schoolchildren aged from 8 to 11 years and those living in rural areas. We conclude that the frequency of multiparasitism is higher in the Nyong et Mfoumou Division. These findings could be helpful in defining and implementing more effective parasitic diseases control strategies.

## Introduction

More than 80% of all living species described to date are parasites **[1]**. They are great in diversity and parasitize a wide range of hosts they often share together. The concomitant presence of two or more parasite species in the same host, called multi-parasitism, appears as the rule than the exception in most biological systems including humans **[1]**.

In infected zones, more than 30% of infections are multiparasitism and this rate can reach up to 80% in some human populations **[1]**. Co-infective parasites interact directly or indirectly through several interspecific mechanisms. These interactions can affect the host’s health because they modify a large number of factors including the host’s susceptibility to other parasites, duration of infection, risks of transmission, clinical signs, therapeutic success and control strategies **[2]**.

Multiparasitism results either from synergistic interactions between two or more parasite species infecting the same host or from a community of risk factors between these parasites, which thus creates statistical associations between them. These factors may be environmental, climatic, related to the host’s behavior and physiological conditions, and the transmission means of these parasites **[2,3]**.

In the Nyong et Mfoumou Division, populations are predominantly rural. They have limited access to safe water, sanitation and basic health services. In addition, climatic and environmental conditions are favorable for the development and persistence of several parasite species. Everywhere in sub-Saharan Africa, school-age children are the most vulnerable segment of the population to parasitic infections, especially intestinal and malaria parasites because of behavioral, hygienic and recreational reasons **[4]**. School-age children infected with intestinal helminths undergo frequent physical and mental sufferings due to anemia, which result in a lack of attention, inability to assimilate knowledge and contribute to absenteeism and school dropouts. Intestinal worms are also responsible for a decreased immunity of children towards malaria **[4]**. In addition, these children are thus most exposed to multiparasitism and its potential consequences **[5]**. Little is known on the frequency and intensity of multiparasitism in Cameroon especially in the Akonolinga area. We therefore conducted a study with the aim to determine the frequency of multiparasitism, identify its determinants in Nyong et Mfoumou Division in the Centre Region of Cameroon and study parasitic associations among species.

## Material and Methods

### Study Area

We conducted a cross-sectional study from September 2017 to July 2018 in Akonolinga, the capital of Nyong et Mfoumou Division in the Centre Region of Cameroon. This division covers an area of 4,300 Km^2^ approximately with a bit more than 100,000 inhabitants **[6]**. The climate is typically equatorial with two discontinuous dry and wet seasons. The annual average rainfall is 2000 mm with an annual average temperature of 24°C **[7]**. The hydrographic network is dense with two main rivers: Nyong and Mfoumou. Several economic activities are developed consisting mainly of agriculture, livestock, fishing, hunting and small businesses. Houses are built in semi-dur with crevices and open joints serving as hideouts for mosquitoes. These villages lack access to potable water. Toilet facilities, made up essentially of pit latrines, are in general poorly constructed and insufficient for the members of a household.

Several prospecting trips were organized on the study area. We randomly selected five (05) Government schools: 3 in rural areas and 2 in urban areas for participant’s recruitment.

### Ethical considerations

This study was approved by the National Ethical Committee of Research for Human Health (Ethical Clearance N°: 2018/01/968/CE/CNERSH/SP) and the Direction of Yaoundé University Teaching Hospital (Research Authorization N°: 894/AR/CHUY/DG/DGA/DMT). However, written informed consent was obtained from parents or legal guardians of all children prior to their inclusion in the study.

### Sample collection

Before sample collection, socio-demographic information of each child data and school environment related information were collected. Afterwards, each child was given a sterile and labeled stool container as well as instructions for the adequate collection of their stools. After collection, stool samples were fixed *in situ* with formalin solution diluted to 10%. Blood samples were collected from each child by pricking finger. Three drops of blood were collected to realize two calibrated thick and one thin blood films which were then air-dried, stored in slide boxes and transported to the Laboratory of Parasitology, Mycology and Parasitic Immunology of the Yaoundé University Teaching Hospital for parasitological examinations.

### Parasitological examinations

Each stool sample was screened by direct examination and Formalin-ether concentration technique **[8]** for the presence of protozoa cysts and helminths eggs and larva. Calibrated thick and thin blood films were stained with May-Grünwald-Giemsa **[9]** and examined for the presence of *Plasmodium* spp and filarial blood stages. All slides were read using the CyScope^®^ microscope (Partec-Sysmex GmbH, Görlitz, Germany) in a blind manner by two qualified technicians. In case of discrepancy, a third qualified technician was called to read the quarreled slides.

### Statistical Analysis

Data was keyed in a Microsoft Excels 2007 spreadsheet then exported to SPSS 16.0 (SPSS, Chicago, Inc., IL, USA) software for statistical analysis. Frequencies of socio-demographic data of participants, and the presence of parasites species were determined. To compare single parasite infections by gender, age groups and living areas, χ^2^–test or Fisher’s exact test were used appropriately. The frequency of multiparasitism was assessed and stratified by gender, age groups and living areas. Multivariable logistic regression was used to investigate associations between parasites and socio-demographic data. *P-value* less than 0.05 were considered statistically significant.

## Results

### Study Population

In total, 416 schoolchildren, 209 from rural area and 207 from urban area, were included in the study. Females accounted for 54.33% (n = 226) of all participants giving a male-to-female sex ratio of 0.8. In addition, 117 and 109 females were reported in rural and urban areas respectively. The age of children ranged from 4 to 15 years with a mean value of 9.17 ± 0.27 years. Children were grouped into 3 age sub-groups of 4 years interval (Table 1). Gender-distribution was similar with respect to these three age groups (χ^2^ = 1.49; df = 2; *P* = 0.47). Children aged from 4 to 7 years and from 12 to 15 years were more frequent in urban areas with 68 and 76 children against 67 and 36 children in rural areas respectively. This pattern was inverted among those aged between 8-11 years where they were more frequent in rural areas than in urban ones (106 versus 63).

**Table 1:** General characteristics of the study population.

|  | Total<br>population<br>(N = 416) | Rural area (n = 209) |  |  | Urban area (n=207) |  |
| --- | --- | --- | --- | --- | --- | --- |
|  |  | Essang-Ndibi<br>(n=73) | Kpwele<br>(n=72) | Eboa<br>(n=64) | EPA<br>(n=76) | Loum<br>(n=131) |
| **Gender %(n)** |  |  |  |  |  |  |
| Male | 45.67 (190) | 45.21 (33) | 50.00 (36) | 35.94 (23) | 38.16 (29) | 52.67 (69) |
| Female | 54.33 (226) | 54.79 (40) | 50.00 (36) | 64.06 (41) | 61.84 (47) | 47.33 (62) |
| **Age sub-group**<br>**(years) %(n)** |  |  |  |  |  |  |
| 4 – 7 | 32.45 (135) | 41.09 (30) | 26.39 (19) | 28.12 (18) | 32.89 (25) | 32.82 (43) |
| 8 – 11 | 40.63 (169) | 34.25 (25) | 62.50 (45) | 56.25 (36) | 42.11 (32) | 23.67 (31) |
| 12 – 15 | 26.92 (112) | 24.66 (18) | 11.11 (8) | 15.63 (10) | 25.00 (19) | 43.51 (57) |

N: Total population, n: sub-population

### Frequencies of parasite species among pupils

Out of the 416 samples examined, 309 (74.28%) were infected with at least one parasite species. Thirteen (13) different parasite taxa including 3 hemoparasites and 10 intestinal parasites (6 protozoa and 4 helminths) were recorded. One hundred and sixty-seven (167) individuals were infected with hemoparasites (40.14%) and 250 (60.10%) with intestinal parasites. Table 2 below summarizes frequencies of different group and parasite species related to total population with regard to age groups, living area and gender. Pupils living in rural areas were more infected with hemoparasites (*P* = 0.0047) and intestinal parasites (*P* = 0.0000) than those living in urban areas.

**Table 2:** Frequency of different parasite groups and species related to Total population, age groups, living area and gender.

| Groups of parasites | Total<br>Population N<br>(%) | Age groups |  |  | *P* | Area |  | *P* | Gender |  | *P* |
| --- | --- | --- | --- | --- | --- | --- | --- | --- | --- | --- | --- |
|  |  | 4 – 7<br>n(%) | 8 – 11<br>n(%) | 12 – 15<br>n(%) |  | Rural<br>n(%) | Urban<br>n(%) |  | M<br>n(%) | F<br>n(%) |  |
| **Hemoparasites** | 167 (40.14) | 46 (34.07) | 77 (45.56) | 44 (39.28) | 0.1244 | 98 (46.88) | 69 (33.33) | 0.0047* | 86(45.26) | 81 (35.84) | 0.3426 |
| *P. falciparum* | 155 (37.26) | 42 (31.11) | 72 (42.60) | 41 (36.60) | 0.1184 | 90 (43.06) | 65 (31.40) | 0.0139* | 73 (38.42) | 82 (36.28) | 0.6532 |
| *M. perstans* | 18 (4.32) | 07 (5.18) | 06 (3.55) | 05 (4.46) | 0.7821 | 13 (6.22) | 05 (2.41) | 0.0565 | 12 (6.31) | 06 (2.65) | 0.0675 |
| *L. loa* | 02 (0.48) | 00 (0.00) | 02 (1.18) | 00 (0.00) | / | 02 (0.95) | 00 (0.00) | / | 01 (0.52) | 01 (0.44) | / |
| **Intestinal Parasites** | 250 (60.10) | 74 (54.81) | 106 (62.72) | 70 (62.50) | 0.3126 | 153 (73.20) | 97 (46.85) | 0.0000* | 152 (80.00) | 98 (43.36) | 0.0011* |
| **Protozoa** | 199 (47.83) | 57 (42.22) | 86 (50.88) | 56 (50.00) | 0.28 | 116 (55.50) | 83 (40.09) | 0.001* | 80 (42.10) | 119 (52.65) | 0.031* |
| *E. coli* | 122 (29.32) | 42 (31.11) | 51 (30.17) | 29 (25.89) | 0.6365 | 89 (42.58) | 33 (15.94) | 0.0001* | 48 (25.26) | 74 (32.74) | 0.095 |
| *E. histolytica* | 99 (23.79) | 22 (16.29) | 45 (26.62) | 32 (28.57) | 0.042* | 46 (22.00) | 53 (25.60) | 0.389 | 38 (20.00) | 61 (26.99) | 0.095 |
| *G. intestinalis* | 17 (4.08) | 04 (2.96) | 10 (5.91) | 03 (2.67) | 0.2943 | 13 (6.22) | 04 (1.93) | 0.0272* | 13 (6.84) | 04 (1.76) | 0.027* |
| *B. hominis* | 06 (1.44) | 03 (2.22) | 01 (0.59) | 02 (1.78) | 0.4652 | 06 (2.87) | 00 (0.00) | 0.014* | 02 (1.05) | 04 (1.76) | 0.541 |
| *Em. intestinalis* | 01 (0.24) | 01 (0.74) | 00 (0.00) | 00 (0.00) | / | 01 (0.47) | 00 (0.00) | / | 01 (0.52) | 00 (0.00) | / |
| *En. nanus* | 13 (3.12) | 06 (4.44) | 04 (2.36) | 03 (2.67) | 0.5568 | 08 (3.82) | 05 (2.41) | 0.407 | 07 (3.68) | 06 (2.65) | 0.547 |
| **Helminths** | 135 (32.45) | 44 (32.59) | 63 (37.27) | 28 (25.00) | 0.0986 | 107 (51.19) | 28 (13.52) | 0.0000* | 50 (26.31) | 85 (37.61) | 0.018* |
| *A. lumbricoides* | 89 (21.39) | 23 (17.04) | 42 (24.85) | 24 (21.42) | 0.2559 | 70 (33.49) | 19 (9.17) | 0.0000* | 32 (16.84) | 57 (25.22) | 0.0379* |
| *T. trichiura* | 77 (18.50) | 30 (22.22) | 33 (19.52) | 14 (12.50) | 0.1332 | 65 (31.10) | 12 (5.79) | 0.0000* | 30 (15.78) | 47 (20.79) | 0.19 |
| *N. americanus* | 04 (0.96) | 03 (2.22) | 01 (0.59) | 00 (0.00) | / | 02 (0.95) | 02 (0.96) | / | 01 (0.52) | 03 (1.32) | / |
| *H. nana* | 01 (0.24) | 00 (0.00) | 01 (0.59) | 00 (0.00) | / | 00 (0.00) | 01 (0.48) | / | 00 (0.00) | 01 (0.44) | / |

*Plasmodium falciparum* was the most common hemoparasites followed by *Mansonella perstans* and *Loa loa*. More than one third (37.26%) of participants had an infection with *P. falciparum*. This infection was not influenced by age group nor gender but was significantly (*P* = 0.0139) higher in rural areas than in urban areas. The overall prevalence of *M. perstans* was 4.32 % and likewise was not influenced by age group nor gender as *Plasmodium* infection. *Loa loa* was found only in rural areas, with infection prevalence of 0.48%.

The prevalence of intestinal helminths was significantly higher in rural areas (51.19%, *P*< 0.0001) and females (37.61%, *P* = 0.018) compared to urban areas (13.52%) and males (26.31%). The highest infection rate among this group of parasites were found for *Ascaris lumbricoides* (21.39%); and its prevalence was also found to be higher in pupils of rural areas (33.49%, *P*< 0.0001) and females (25.22%, *P* = 0.018). Likewise, a high infection rate was reported for *Trichirus trichiura* with a value of 18.51%. Its prevalence was significantly higher in rural areas than in urban areas (*P*< 0.001); participants’ age did not influence the risk for infection with these both parasites (*P > 0.05*). *Necator americanus* and *Hymenolepis nana* were also found in this study with prevalence of 0.96% and 0.24% respectively (Table 2).

*Entamoeba coli* was the main intestinal protozoa reported in this study (29.33%) and its prevalence was significantly higher in rural areas (42.58% versus 15.9%; *P* = 0.0001). The overall prevalence of *E. histolytica/dispar* was 23.80% and was significantly higher among schoolchildren aged from 12 to 15 years (*P* = 0.042). Prevalence of *Giardia intestinalis*, *Endolimax nanus* and *Blastocystis hominis* were 4.09%, 3.13% and 1.44% respectively. *Embadomonas intestinalis* was found only in rural areas with a prevalence of 0.24% (Table 2).

### Parasitic infra-communities

Amongst 416 schoolchildren recruited, only 124 (29.80%) were infected with only one parasite species and up to 185 (44.47%) were infected with two and more parasites species. The maximum number of parasite species found in a host was 5 and the mean specific richness was 1.43 ± 0.01 per individual. One hundred and eleven (26.68%) harbored two parasites species concurrently (bi-parasitism). There were 47 (11.29%) cases of 3 parasite infra-community (tri-parasitism); 24 (5.76%) cases of 4 parasites infra-community (quadri-parasitism); and 3 (0.72%) cases of 5 parasites infra-community (penta-parasitism). The Table 3 displays different parasitic infra-communities found in the study population.

**Table 3:**
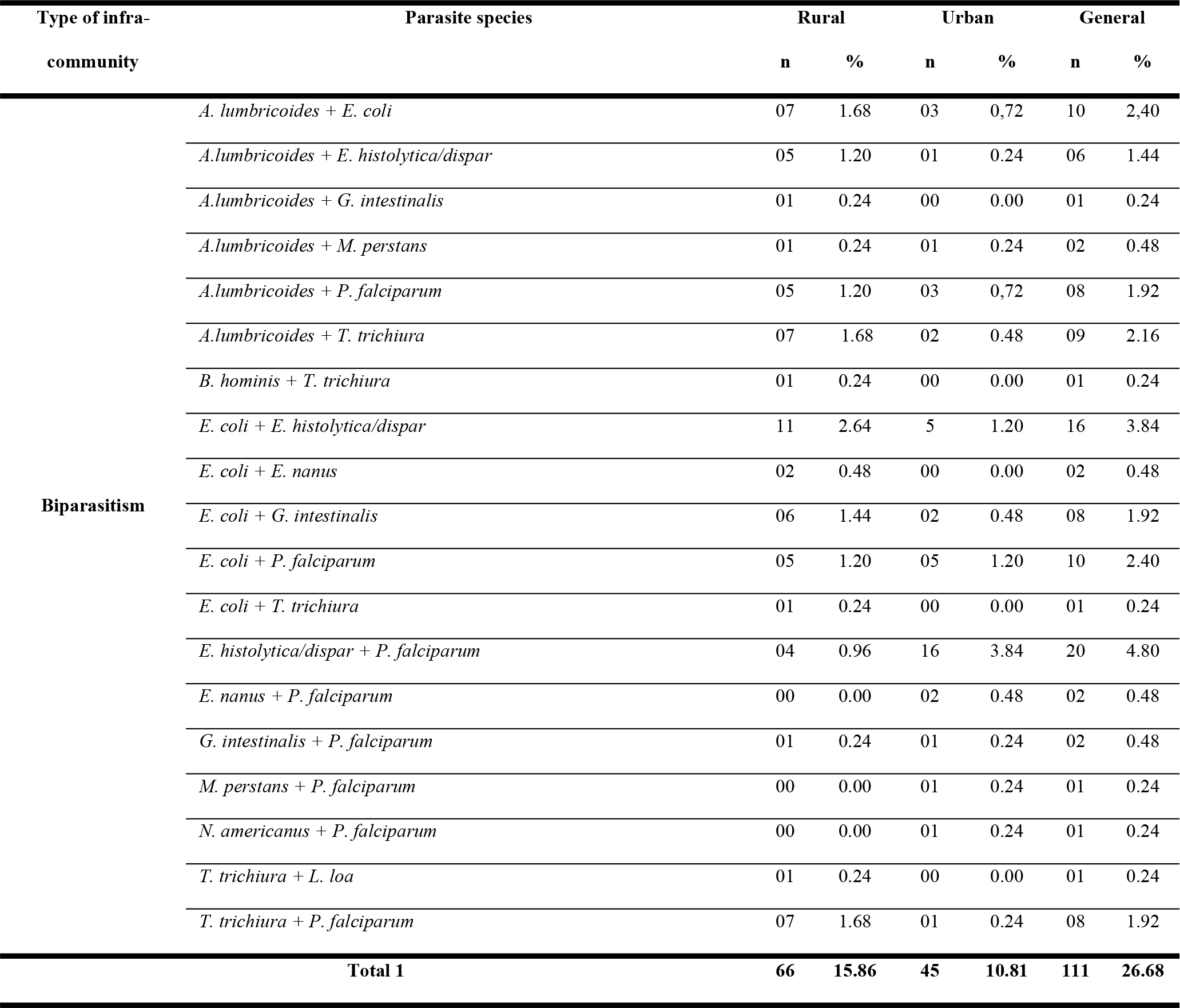
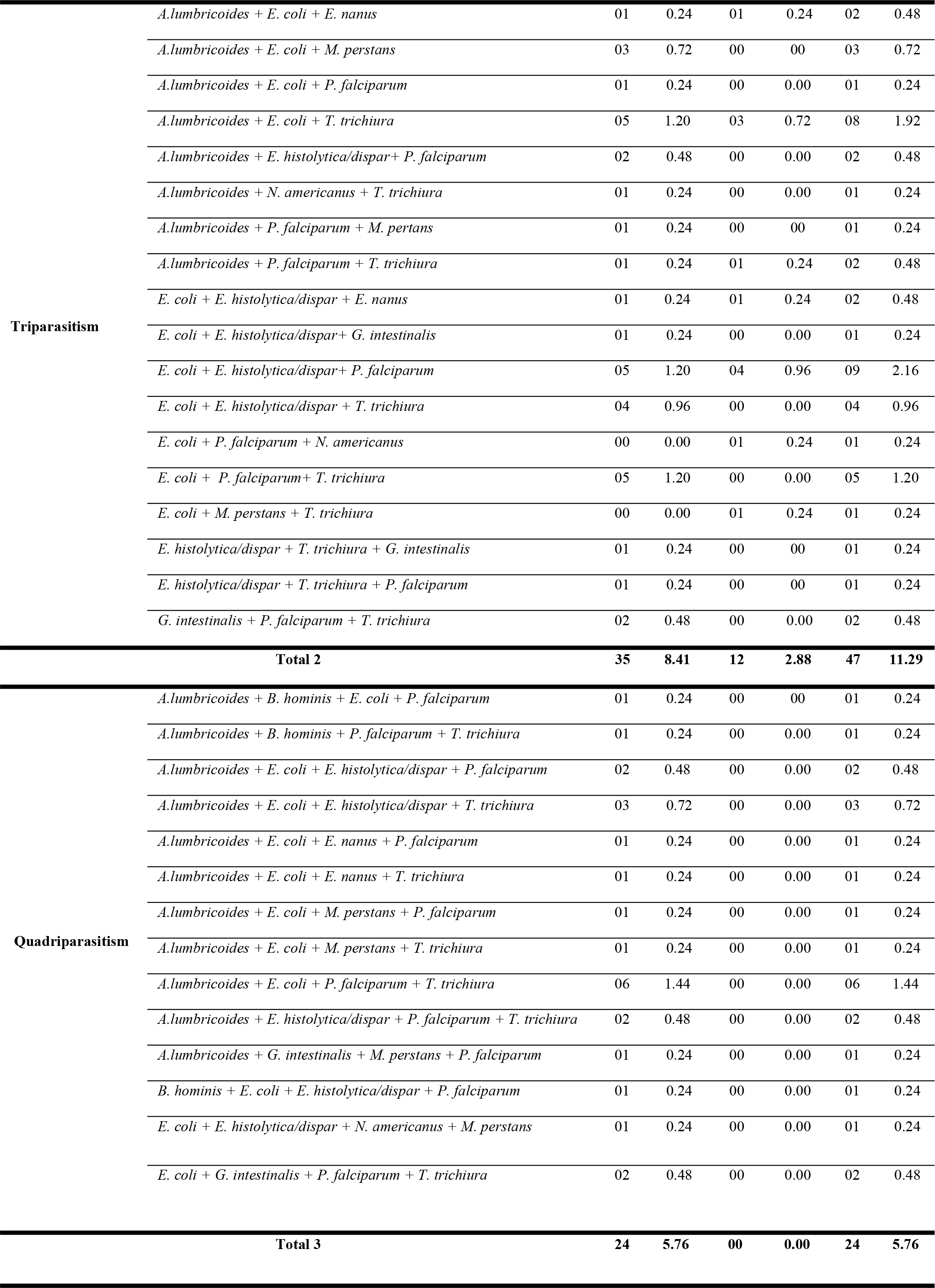
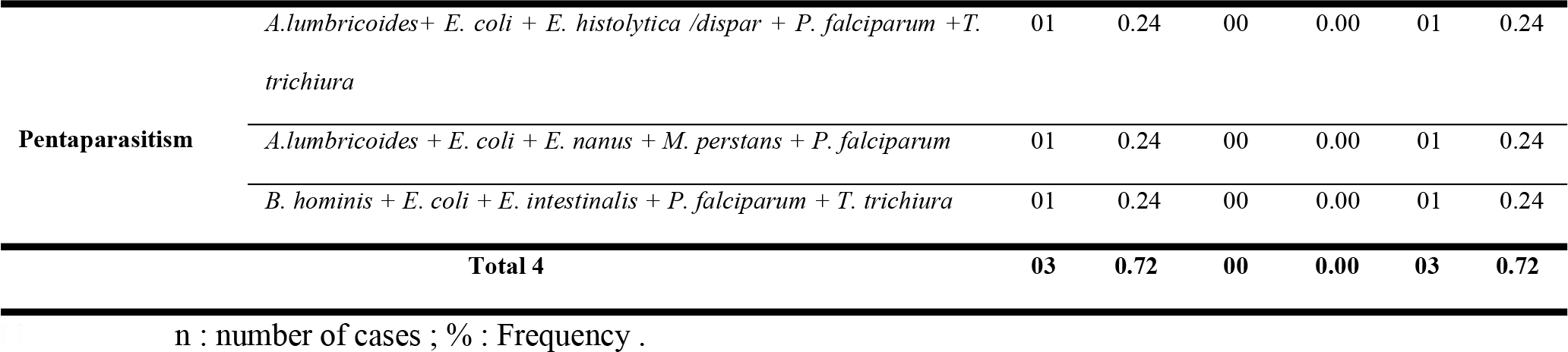
Different parasitic infra-communities observed according to the living area.

### Risks factors of Monoparasitism and Multiparasitism

Table 4 displays the results of the logistic regression analysis which identifies associated factors of monoparasitism and multiparasitism. Age group and living area were associated with a high risk of monoparasitism. In fact, schoolchildren aged from 8 to 11 years were almost twice as likely to be infected with single parasite compared to those aged from 4 to 7 years (OR = 1.92; 95% IC: 1.03-3.59; *P* = 0.0403). Schoolchildren living in urban areas were less likely to be infected with one parasite than those living in rural areas (OR = 0.56; 95% IC: 0.32-0.99; *P* = 0.0492). The risk of multiparasitism was significantly influenced by gender, age group and living areas. In fact, the risk was higher for females (OR = 2/12; 95% IC: 1.26-3.57; *P* = 0.0046) and schoolchildren aged from 8 to 11 years (OR = 1.70; 95% IC: 1.09-3.15; *P* = 0.0089) compared to males and those aged from 4 to 7 years, respectively. In contrast, the risk of multiparasitism was low among schoolchildren living in urban areas (OR = 0.16; 95%IC: 0.09-0.28; *P* = 0.0001) compared to those living in rural areas.

**Table 4:** Associated variables of risk of Monoparasitism and multiple Multiparasitism.

| Monoparasitism |  |  |  |  | Multiparasitism |  |  |  |
| --- | --- | --- | --- | --- | --- | --- | --- | --- |
| Variables | Brut OR<br>[IC95%] | P | Adjusted OR<br>[IC95%] | P | Brut OR<br>[IC95%] | P | Adjusted OR<br>[IC95%] | P |
| **Gender** |  |  |  |  |  |  |  |  |
| Male | 1 |  | 1 |  | 1 |  | 1 |  |
| Female | 1,23<br>[0,73 - 2,06] | 0,4382 | 1,26<br>[0,74 - 2,13] | 0,3904 | 1,99<br>[1,22 - 3,22] | 0,0055* | 2,12<br>[1,26 - 3,57] | 0,0046* |
| **Age Group** |  |  |  |  |  |  |  |  |
| 4 - 7 ans | 1 |  | 1 |  | 1 |  | 1 |  |
| 8 - 11 ans | 2,03<br>[1,10 - 3,77] | 0,0243* | 1,92<br>[1,03 - 3,59] | 0,0403* | 1,98<br>[1,12 - 3,50] | 0,0192* | 1,70<br>[1,09 - 3,15] | 0,0089* |
| 12 - 15 ans | 1,20<br>[0,62 - 2,32] | 0,5895 | 1,35<br>[0,69 - 2,65] | 0,3805 | 1,24<br>[0,68 - 2,25] | 0,4901 | 1,91<br>[0,99 - 3,71] | 0,0551 |
| **Living area** |  |  |  |  |  |  |  |  |
| Rural | 1 |  | 1 |  | 1 |  | 1 |  |
| Urban | 0,53<br>[0,30 - 0,92] | 0,0246* | 0,56<br>[0,32 - 0,99] | 0,0492* | 0,17<br>[0,10 - 0,28] | <<br>0,0001* | 0,16<br>[0,09 - 0,28] | <<br>0,0001* |

OR: Odd Ratio; 95%CI: 95% Confidence Interval; P: P-value; (\*): Significant

### Parasitic associations

All significant association (*P*< 0.05) between parasites, gender, age groups and living areas resulting from multivariable logistic regression are summarized in Table 5. *Trichuris trichiura* showed a significant and positive association with *A. lumbricoides* (aOR = 2.49; 95%CI = 1.39-4.43) and *E. coli* (aOR = 2.95; 95%CI = 0.17-5.13) but a significant and negative association with the urban area. A positive association was found between *M. perstans* and *A. lumbricoides* (aOR = 1.78; 95% CI = 0.62 −5.09) but a negative association with the urban setting (aOR = 0.32; 95% CI = 0.11-0.94). *E. histolytica/dispar* showed a positive association with *T. trichiura* (aOR = 2.10; 95% CI = 1.04-4.20), *E. coli* (aOR = 4.35; 95%CI = 2.61 - 7.25) and the age group between 8 and 11 years (aOR = 2.06; 95%CI = 1.11 −3.85). A significant and negative association was found between *P. falciparum* and the urban environment (aOR = 0.62; 95%CI = 0.41-0.93).

**Table 5:** Association between a particular parasitic and gender, age group and any remaining parasites.

| Parasites | Co-variables | aOR (95%IC) | P |
| --- | --- | --- | --- |
| *T. trichiura* | *A. lumbricoides* | 2,49 (1,39-4,43) | <0,001 |
|  | *E. coli* | 2,95 (0,17-5,13) | <0,001 |
|  | Rural area | 0,14 (0,07 - 0,26) | < 0,001 |
| *A. lumbricoides* | *E. coli* | 4,95 (2,87 – 8,48) | 0,011 |
|  | *M. perstans* | 1,79 (0,63 – 5,09) | 0,001 |
|  | Female | 1,71 (1,03 - 2,85) | 0,04 |
|  | Rural area | 0,19 (0,10 - 0,33) | < 0,0001 |
| *M. perstans* | Rural area | 0,32 (0,11 - 0,94) | 0,04 |
| *E. histolytica/dispar* | *T. trichiura* | 2,10 (1,04 – 4,20) | 0,036 |
|  | *E. coli* | 4,35 (2,61 – 7,25) | 0,001 |
|  | [8 -11[ years | 2,06 (1,11 - 3,85) | 0,02 |
| *E. coli* | Rural area | 0,24 (0,15 - 0,40) | < 0,001 |
| *P. falciparum* | Rural area | 0,62 (0,41 - 0,93) | 0,02 |

Adjusted OR: Odd Ratio; 95%CI: 95% Confidence Interval; P: P-value.

## Discussion

The study showed that the frequency of multiparasitism is higher than that of monoparasitism in Akonolinga, Nyong et Mfoumou Division, Centre Region of Cameroon. Approximately ¾ of the population studied were infected with at least one parasitic species. This infection rate is lower than that found by Kimbi *et al*. **[10]** more than 82%, among schoolchildren in the South-West Region of Cameroon. It is similar to that of Zeukeng *et al*. **[7]**77.2% among general population in the Centre Region of the same country. Conversely, our finding is higher than 26.6% obtained by Lehman *et al*. **[11]**, and 8.5% by Khan Payne *et al*. **[12]** in Littoral, Centre and West regions of Cameroon respectively. This shows that despite the fact that schoolchildren are the main target of parasitic infection control strategies, they are still the segment of the population most vulnerable to parasitic infections. The highest prevalence was found in schoolchildren of rural areas. As we noticed in the rural area, drinking unsafe water, wearing open shoes and using latrines, if they existed, inappropriately maintained by children could justify these results.

A total of 13 parasitic species were found in this study population. Ten of them were also reported by M’bondoukwe *et al*. **[13]** in Gabon (*P. falciparum*, *L. loa*, *M. perstans*, *G. intestinalis*, *E. coli*, *E. histolytica/dispar*, *B. hominis*, *A. lumbricoides*, *T. trichiura* and *N. americanus*). Similarly, Raso *et al*. **[14]** and Coulibaly *et al*. **[15]** reported both in Côte d’Ivoire, nine and eight parasite species found in our study (*N. americanus*, *T. trichiura*, *A. lumbricoides*, *E. coli*, *E. histolytica*/*dispar*, *B. hominis*, *E. nana*, *G. intestinalis* and *P. falciparum*). In the same country were found 8 species of intestinal parasites out of the 10 obtained in our study. These results confirm that in sub-Saharan Africa, environmental and climatic conditions are favorable for the development and persistence of several parasite species.

*Plasmodium falciparum* being the only species of that genus found and this study confirms that it is the main malarial agent in Cameroon as previously reported by Kimbi *et al*. **[10]** and Payne Khan *et al*. **[12]**. Malaria prevalence was higher in the rural area; this is consistent with findings of Olurongbe *et al*. **[16]** and Kimbi *et al*. **[10]** and may be due to higher risk of contact with mosquito vectors as a result of a higher presence of their breeding sites, lower level of education and lower rate of preventive methods against malaria in this area compared with urban ones **[10]**.

*Entamoeba coli* (29.33%) and *E. histolytica/dispar* (23.80%) were the commonest intestinal protozoa found in this study. The prevalence of the former protozoa is similar to that reported by Coulibaly *et al*. **[15]** in Côte d’Ivoire (31.8%) but higher than that obtained by M’bondoukwé *et al*. **[13]** in Gabon (22.2%). As regards *E. histolytica/dispar*, its overall prevalence was higher than those obtained by the above mentioned authors who had values of 7.4 % and 9.3% respectively. The higher prevalence of amoebae in this study could be justified by the fact that study period coincided with the harvest and increased consumption of fruits such as mangoes. It should be noticed that a large proportion of children, especially in rural areas, were consuming fruits without prior washing them and/or washing their hands.

*Ascaris lumbricoides* (21.4%) and *T. trichiura* (18.5%) were the main parasitic helminths found in this study. High prevalence of *A. lumbricoides* and *T. trichiura* observed among intestinal helminths could be explained by the fact that both are fecal-orally transmitted and their epidemiology dependent on individual and community hygienic habits and human waste disposal methods and then, subsequently affect the level of environmental contamination. Ova from both species are equipped with outer coat that enables them to resist adverse external environmental conditions and enhances their survival and higher probability of transmission **[17]**. Besides, the prevalence of both soil transmitted helminths were higher than those reported by Khan Payne *et al*. **[12]** in the West Region of Cameroon (*A. lumbricoides* 4% and *T. trichiura* 4.1%) and M’Bondoukwé *et al*. **[13]** in Gabon (*A. lumbricoides* 13.7% and *T. trichiura* 11.8%). However, these findings are lower than those found by Kimbi *et al*. **[10]** in the South West Region of Cameroon (*A. lumbricoides* 30.21% and *T. trichiura* 25.98%) and Ruto *and* Mulambalah **[17]**in Kenya (*A. lumbricoides* 55.8% and *T. trichiura* 26.9%).

Approximately 45% of the study population was infected with two or more parasites. The maximum number of parasitic taxa found in a simple host was 5 and the mean specific richness was 1.43 ± 0.01 parasite per individual. Our finding agrees with Tchuem Tchuenté *et al*. **[18]** and Kimbi *et al*. **[10]** in Cameroon; Raso *et al*. **[14]** and Hürlimann *et al*. **[19]** in Côte d’Ivoire; Ruto *and* Mulambalah **[17]** in Kenya. These findings are in line with the statement of Petney and Andrews **[1]** on the fact that: “multiparasitism is the rule rather than the exception in most biological systems and the co-infection rate can reach 80% in some human populations”. The biparasitism was the main parasitic association in this study with the association *P. falciparum* + *E. histolytica/dispar* primarily reported (18%) among children. This can be attributed to the high prevalence rates and local endemicity of these parasites in the area.

Statistically significant associations were found between some parasitic taxa especially between *A.lumbricoides* and *T. trichiura* and between *E. coli* and *E. histolytica/dispar*. Such observations were also previously reported by Raso *et al*. **[14]**, Coulibaly *et al*. **[15]** and Hürlimann *et al*. **[19]**. These parasitic taxa share the same routes of transmission to humans through ingestion of contaminated food or drinking water with parasite infesting development stages. The lack of hygiene and poor health conditions can also favor the transmission of these parasites. The significant association between *A. lumbricoides* and *M. perstans*, two helminths with different ecological niches in the host, would mean a synergy between these parasites via host immunity. Indeed, all helminthiasis are chronic infections hallmarked by a strong immune response to Th2 cell-mediated dominated by an increase in anti-inflammatory cytokines **[20]**. Thus, the installation of a helminth would create conditions favorable to the installation of other helminths within the host.

## Conclusion

Our study pointed out that parasitic infections are highly prevalent in Akonolinga especially *P. falciparum*, *Ascaris lumbricoides*, *Entamoeba coli* and *Entamoeba histolytica/dispar* infections. The frequency of multiparasitism is higher than that of monoparasitism in Akonolinga, Nyong et Mfoumou Division. Among parasitic infra-communities found, bi-parasitism was more frequent. The study also outlined many parasitic statistical associations. These findings could be helpful in defining and implementing more effective parasitic disease control strategies in the Nyong et Mfoumou Division.

## Acknowledgments

The authors are grateful to all pupils who voluntarily accepted to participate in this study as well as their parents/guardians. The authors would like to acknowledge the valuable assistance of teachers and officials of schools where children were recruited. The technical assistance of the staff of the Laboratory of Parasitology, Mycology and Parasite Immunology of CHUY is greatly appreciated. Finally, we thank Prof. **SAME EKOBO Albert** who gave us precious advice for the study design.

## Supporting Information Legends

**S1 Checklist: STROBE Checklist**

**S1 Table**. **General characteristics of the study population.** N: Total population, n: sub-population

**S2 Table. Frequency of different parasite groups and species related to Total population, age groups, living area and gender.** n: number of positive; %: Frequency; *P*: P-value; **\***: Statistically significant at P-value less than 0.05; P: *Plasmodium*; M: *Mansonella*; L: *Loa*; E: *Entamoeba*; G: *Giardia*; A: *Ascaris*; T: *Trichuris*; N: *Necator*; H: *Hymenolepis*; Em: *Embadomonas*; B: *Blastocystis*; En: *Endolimax*

**S3 Table 3. Different parasitic infra-communities observed according to the living area.** n: number of cases; %: Frequency

**S4 Table 4. Associated variables of risk of Monoparasitism and multiple Multiparasitism.** OR: Odd Ratio; 95%CI: 95% Confidence Interval; P: P-value; (*): Significant

**S5 Table 5. Association between a particular parasitic and gender, age group and any remaining parasites.** Adjusted OR: Odd Ratio; 95%CI: 95% Confidence Interval; *P*: *P*-value.

## References

1. Petney TN, Andrews RH (1988)Multiparasite communities in animals and humans: frequency, structure and pathogenic significance. International Journal for Parasitology 28:377–393.

2. Vaumourin E, Vourch G, Gasqui P, Vayssier-Taussat M (2015)The importance of multiparasitism: examining the consequences of co-infection for human and animal health. Parasites & Vectors 8: 545. Doi 10.1186/s13071-015-1167-9.

3. Lello J, Boag B, Fenton A, Stevenson IR, Hudson PJ (2013)Competition and mutualism among the gut helminthes on a mammalian host. Nature 428:840–844.

4. Hamit MA, Tidjani A, Otchom BB, Tidjani MT, BilongBilong CF (2013) An Epidemiological Assessment of the Infectious forms of Intestinal Helminths in School Children from Chad. Journal of Biology and Life Science 4:2.

5. Brooker S, Clements ACA, Hotez PJ, Hay SI, Tatem AJ, Bundy DAP, et al. (2006) The co-distribution of *Plasmodium falciparum* and hookworm among African schoolchildren. Malaria Journal 5:99. doi:10.1186/1475-2875-5-99.

6. Bayaga HN, Guedje NM, Biye EH (2017) Approche enthobotanique et etnopharmacologique des plantes utilisées dans le traitement traditionnel de l’ulcère de Buruli à Akonolinga (Cameroun). International Journal of Biological and Chemical Sciences 11(4):1523–1541.

7. Zeukeng F, Tchinda VHM, Bigoga JD, Seumen CHT, Ndzi ES, Abonweh G, et al. (2014)Co-infections of malaria and geohelminthiasis in two rural communities of Nkassomo and Vian in the Mfou Health District, Cameroon. PLoS Neglected Tropical Diseases 8(10):e3236.

8. Uga S, Tanaka K, Iwamoto N (2010) Evaluation and modification of the formalin-ether sedimentation technique. Tropical Biomedecine 27(2): 177–184.

9. World Health Organization (2010) Basic Malaria Microscopy. Part I. Learner’s Guide, Second Edition.

10. Kimbi HL, Lum E, Wanji S, Mbuh JV, Ndamukong-Nyanga JL, EyongEbanga EJ, et al. (2012) Co-infections of Asymptomatic Malaria and Soil-Transmitted Helminths in School Children in Localities with Different Levels of Urbanisation in the Mount Cameroon Region. Journal of Bacteriology & Parasitology 3:2.

11. Lehman LG, Kouadjip Nono L, BilongBilong CF(2012)Diagnostic des parasitoses intestinalis à l’aide de la microscopie à fluorescence. Médécine d’Afrique Noire 59: 7.

12. Khan Payne V, Lontua FR, Ngangnang GR, Megwi L, Mbong E, Yamssi C, et al. (2017) Prevalence and Intensity of infection of gastro-intestinal parasites in Babadjou, west of Cameroon. International Journal of Clinical and Experimental Medical Sciences 3(2):14–22.

13. M’bondoukwé NP, Kendjo E, Mawili-Mboumba DP, Lengongo KJV, Mbouoronde OC, Nkoghe D, et al. (2018)Prevalence of and risk factors for malaria, filariasis, and intestinal parasites as single infections or co-infections in different settlements of Gabon, Central Africa. Infectious Diseases of Poverty 7:6.

14. Rosa G, Luginbühl A, Adjoua CA, Tian-Bi TN, Silué DK, Matthys B, et al. (2004)Multiple parasite infections and their relationship to self-reported morbidity in a community of rural Côte d’Ivoire. International Journal of Epidemiology 33:1092–1102.

15. Coulibaly JT, Fürst T, Silué KD, SKnopp S, Hauri D, Ouattara M, et al. (2012) Intestinal parasitic infections in schoolchildren in different settings of Côte d’Ivoire: effect of diagnostic approach and implications for control. Parasites & Vectors 5:135.

16. Olurongbe O, Adegbayi AM, Bolaji OS, Akindele AA, Adefioye AO(2011)Asymptomatic falciparum malaria and helminth co-infection among school children in Osogbo, Nigeria. J Res Med Sci 16:680–686.

17. Ruto J, Mulambalah CS (2016). Epidemiology of parasitism and poly-parasitism involving intestinal helminths among school children from different residential settings in Nandi County, Kenya. CHRISMED J Health Res 3:168–72.

18. TchuemTchuenté LA, Behnke JM, Gilbert FS, Southgate VR, Vercruysse J, (2004)Polyparasitism with Schistosoma haematobium and soil-transmitted helminth infections among school children in Loum, Cameroon. Tropical Medecine and International Health 8(11): 975–986.

19. Hürlimann E., Yapi R. B., Houngbedji C. A., Schmidlin T., Kouadio B. A., Silué K. D. et al. (2014). The epidemiology of polyparasitism and implications for morbidity in two rural communities of Côte d’Ivoire. Parasites & Vectors 7:81.

20. Bwanika R., Kato D.C., Welishe J., Mwandah C.D. (2018). Cytokine profiles among patients co‑infected with Plasmodium falciparum malaria and soil borne helminths attending Kampala International University Teaching Hospital, in Uganda. Allergy, Asthma & Clinical Immunology 14:10.

